# SHIFTS IN THE CONFLICT-COEXISTENCE CONTINUUM: EXPLORING SOCIAL-ECOLOGICAL DETERMINANTS OF HUMAN-ELEPHANT INTERACTIONS

**DOI:** 10.1101/2022.08.24.505141

**Authors:** Grace S. Malley, L.J. Gorenflo

**Affiliations:** Department of Geography, Pennsylvania State University, University Park, Pennsylvania, United States of America.; Institute of Resource Assessment, University of Dar es Salaam, Dar es Salaam, Tanzania.; Department of Landscape Architecture, Pennsylvania State University, University Park, Pennsylvania, United States of America.

## Abstract

In Morogoro Region of south-central Tanzania, loss of crops and safety concerns due to elephants compromises livelihoods in many rural communities relying on subsistence agriculture. Using a social-ecological system framework to examine conflict-coexistence between people and elephants, this paper explores drivers that influence human-elephant interaction and subsistence farmer attitudes towards elephants in 10 villages from three different districts. Surveys and interviews document experiences interacting with elephants along with direct and indirect costs incurred in sharing the landscape, revealing different tolerance levels by residents of subject communities towards elephants that have important implications for elephant conservation. Rather than uniformly negative beliefs about elephants, analyses reveal that over the past decade a shift has occurred from largely favorable to unfavorable. The variables influencing attitudes included amounts of crops lost to elephants, perceived benefits from elephants, amounts of crops lost to other causes, perceived trend of human-elephant conflict (HEC) in the past three decades and level of education. Villager tolerance varied by level of income, perception on how the community coexists with elephants, amounts of crops lost to elephants and compensation. The study contributes to understanding how HEC is affecting the relationship between people and elephants, revealing a shift in the conflict-coexistence continuum from positive to broadly negative and identifying characteristics underlying varying tolerance towards elephants in different communities. Rather than a static condition, HEC emerges under specific conditions at particular times and places through varying, uneven interactions between rural villagers and elephants. In communities vulnerable to food insecurity, such conflict exacerbates existing problems of poverty, social inequality, and feelings of oppression. Addressing the causes of HEC, when possible, will be essential to elephant conservation as well as to improving the wellbeing of rural villagers.

## Introduction

Currently, more than 1 million species of plants and animals are threatened with extinction(1). Termed “Anthropocene defaunation,” this mass extinction is anthropogenically driven as growing human demand and geographically expanding human presence increasingly place people in competition with wildlife for key resources (2)(3,4)With global human population projected to exceed 11 billion by the end of the 21^st^ century, anthropogenic impacts and associated habitat modifications likely will intensify in coming decades (5). Growing human population and expanding geographic presence often forces wildlife and people into closer contact, frequently leading to adverse impacts on both human wellbeing and biodiversity. One of the most direct impacts is human-wildlife conflict (HWC), a growing challenge in the developing world.

In Sub-Saharan Africa, HWC is an increasingly serious problem as the livelihood of millions of people depends on subsistence agriculture(6,7)(8). Although impacts involve a range of wildlife types, those from large charismatic species are particularly noticeable. Animals with large home ranges amid shrinking natural habitat increasingly experience conflicts with humans whose geographic presence is expanding, often into the same shrinking natural habitat (9,10). One noteworthy example of such HWC is growing conflict between elephants and rural villagers in Tanzania, a country with many elephants and a steadily increasing human population that is expanding into elephant habitat (11). This expansion has led to frequent encounters between people and elephants, often taking the form of crop raiding, property and water facility damage, and human injury. Loss of subsistence crops to elephants increases food insecurity, adding challenges to already vulnerable livelihoods in the face of other stressors such as climate change.

In this paper, we explore the growing tension between people and elephants in the Morogoro Region of south-central Tanzania, focusing on 10 villages in three districts that host subsistence agriculturalists as well as protected areas with large elephant populations. This part of Tanzania recently has experienced land use conflict between smallholder farmers and pastoralists, in conjunction with growing population creating pressure on resources. This pressure becomes more intense when elephants moving between protected areas through wildlife migration corridors encounter people. The result places poor subsistence farmers in direct conflict with a native species important to tourism and categorized as vulnerable to extinction by the International Union for Conservation of Nature’s Red List of Threatened Species (12)

Human-elephant interaction (HEI) occurs along an array of positive to negative experiences that vary in intensity, severity, frequency, and scale (9), with negative experiences known as human-elephant conflict (HEC). To explore HEI, we employ a social-ecological system (SES) framework (13). SES integrates ecosystems and human societies into an interdependent framework, the biophysical factors in the former affecting social factors in the latter, and *vice versa,* through a process of feedback that influences the human-wildlife relationship (14). This paper uses an SES framework to explore social-ecological drivers that influence HEI and subsistence farmer attitudes towards elephants. Lischka et al.(15)proposed a comprehensive framework for understanding SES components in the context of human-wildlife interaction (Figure 1). Organized in tiers of organizational levels from individual to society/ecosystem level, the changing relationship between humans and elephants is reflected in varying human attitudes and behavior over space and time (16).

**Figure 1:**
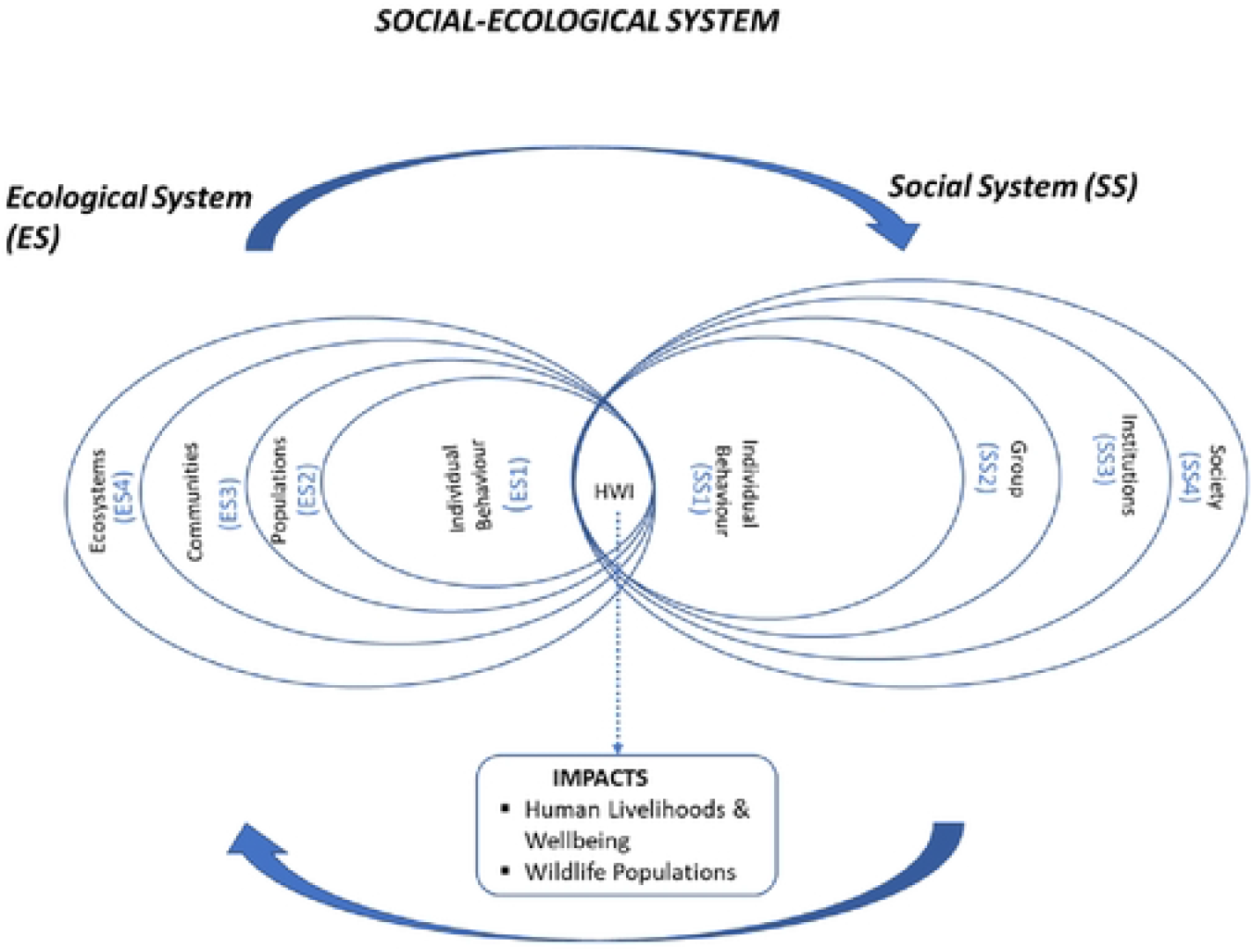
Conceptual framework for analyzing social-ecological systems and human-wildlife interactions (adapted from Lischka ct al.. 2018).

In the context of HEI, an SES consists of two spheres:

i. *Ecological System:* This involves factors associated with elephant ecology and behavior. This will comprise patterns of elephant movement and feeding, environmental preferences, and group dynamics that guide where elephants occur and the activities they pursue. It may also include crop raiding, risk avoidance, adaptation to mitigation measures, demographic characteristics, and bio-physical factors and their influence on the risk of crop raiding. But These factors can vary from individuals to populations and at a local to ecosystem level.
ii. *Social System:* This encompasses cultural, socio-economic, historical, political, knowledge, institutional and individual demographic factors influencing perceptions, values, and behavior of people concerning elephant conservation and the feasibility of different mitigation strategies. Human perceptions can vary considerably based on societal and individual attributes, including values, experiences, emotions, educational levels and wealth (17), (18). Linked to perception are attitudes which are subjective in nature and help determine if HEI is seen as conflict or coexistence. Attitude can be favorable or unfavorable with respect to a particular event or issue (17). Attitudes and perceptions shape the nature of interaction and affect the tolerance of individuals interacting with wildlife, such as the ability to endure the costs of sharing space with wildlife.

Interactions among individuals, groups of people, institutions and societies determine the level and the nature of relationships between people and wildlife, resulting in different outcomes, either desirable or undesirable (19). The relationship between humans and a wildlife species can vary over space and time even for the same individuals or species, depending on the interaction of social and ecological systems. Nyhus (9)proposed a theoretical framework that categorizes HWI (Figure 2). The continuum presents different dimensions of HWI and associated outcomes, either conflict or coexistence, depending on the level of severity along axes. Positive interactions resulting in minor impacts that occur with moderate frequency, such as people and peacocks in Figure 2, suggest high potential for coexistence. Infrequent interactions, negative and severe, such as people and sharks, suggest low potential for coexistence.

**Figure 2:**
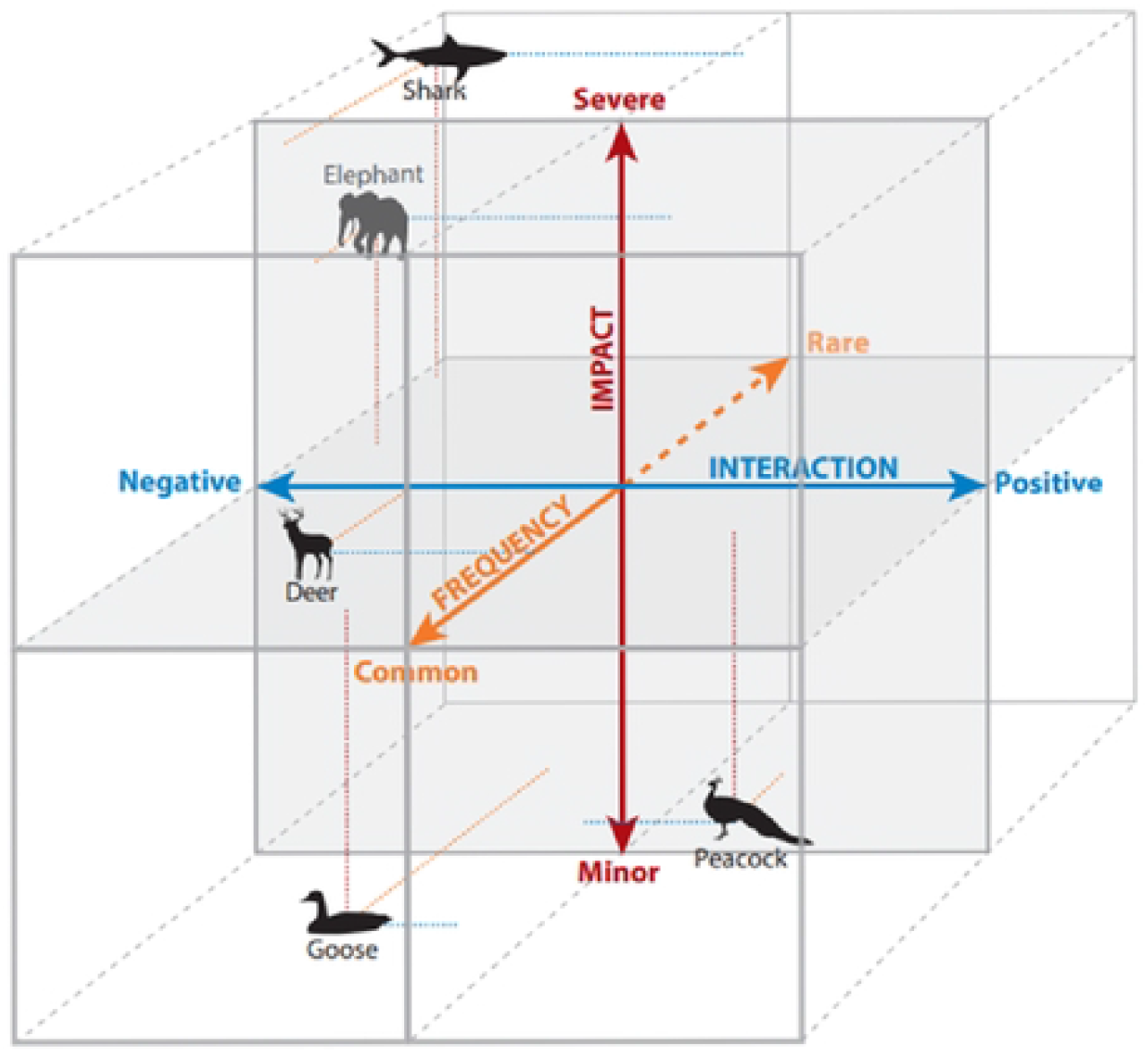
Human-wildlife conflict-coexistence continuum (Nyhus, 2016.)

In the following pages we explore social-ecological components that influence human-elephant relationships in selected Morogoro Region villages. The study examines shifts along the conflict-coexistence continuum—from positive (coexistence) to negative (conflicts). Although previous studies in parts of the study area examined the HEI (20), (21), (22), little is known about how HEI has varied or about underlying SES drivers in the landscape. The primary objective of this study is to provide this additional understanding, ultimately to help design contextualized mitigation measures.

## Materials and Methods

### Study Area

The study focuses on three districts in Morogoro Region in south-central Tanzania—Kilombero, Morogoro Rural, and Mvomero (Figure 3). The districts host portions of Nyerere National Park, Udzungwa Mountains National Park (UMNP), Mikumi National Park and Wami-Mbiki Wildlife Management Area (WMA). Wami-Mbiki WMA acts like a steppingstone between the Southern and Northern Tanzania elephant populations (23), and all three national parks contain substantial elephant populations (24). According to Tanzania Wildlife Authority (TAWA) records, in 2020 Kilombero, Morogoro Rural, and Mvomero districts logged some of the highest incidents of HEC in Tanzania. For example, the records for Mvomero and Kilombero districts showed a total of 352 and 389 crop raiding incidents, respectively, for 2020 alone. The human population and associated activities in this area have been growing in recent decades, providing a possible explanation for such high levels of HEC in these districts. In Kilombero District, the population increased from 321,611 in 2002 to 407,880 in 2012, growing at an average annual rate of 3.7 percent (25). In Mvomero District, human population grew at an average rate of 2.6 percent between 2002 and 2012, increasing from 259,347 to 312,109. Morogoro Rural District also experienced population growth, though a more modest increase from 263, 920 in 2002 to 286,248 people in 2012. Immigration for agriculture and pastoralism plays a major role in the population increase in all three districts (26).

**Figure 3:**
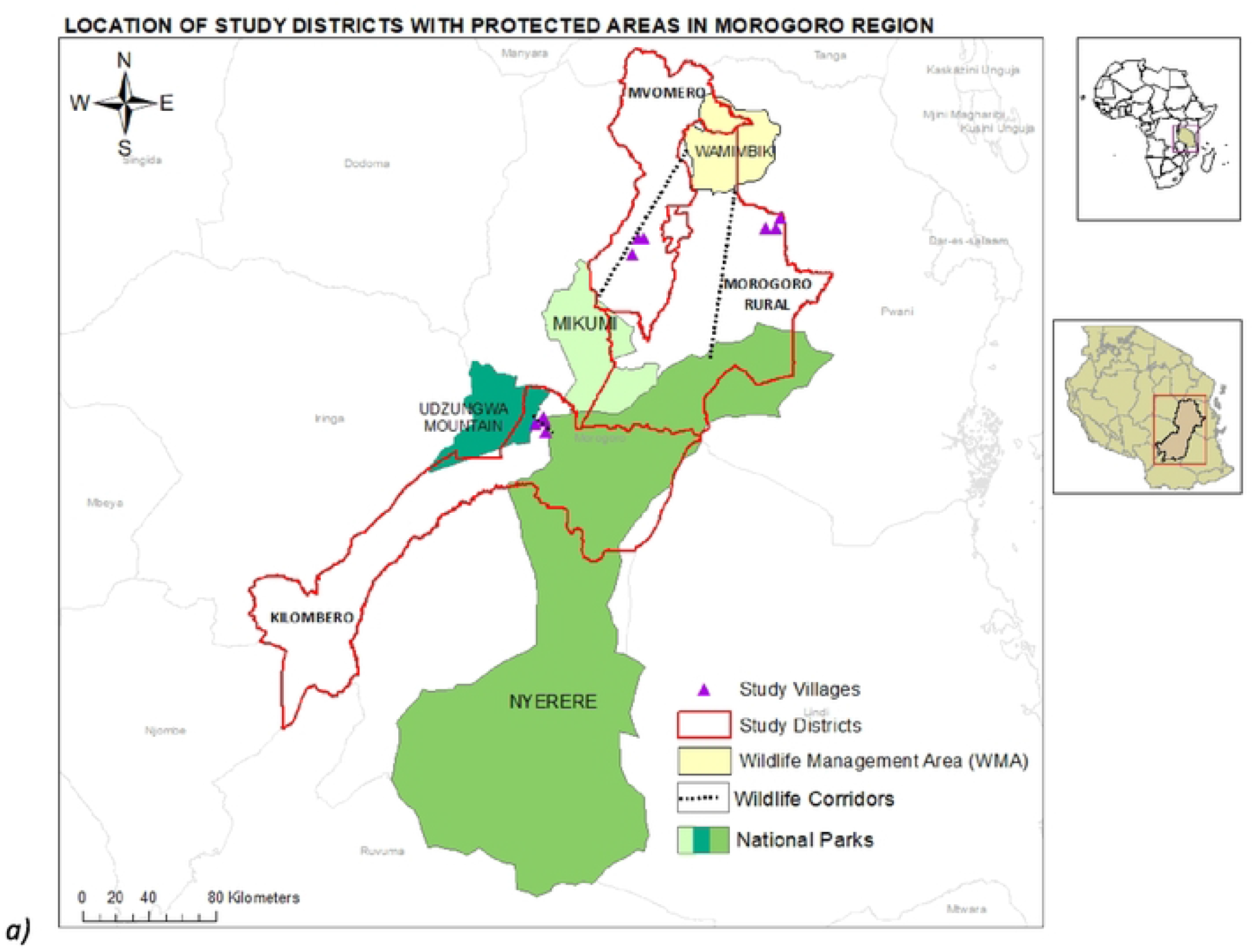
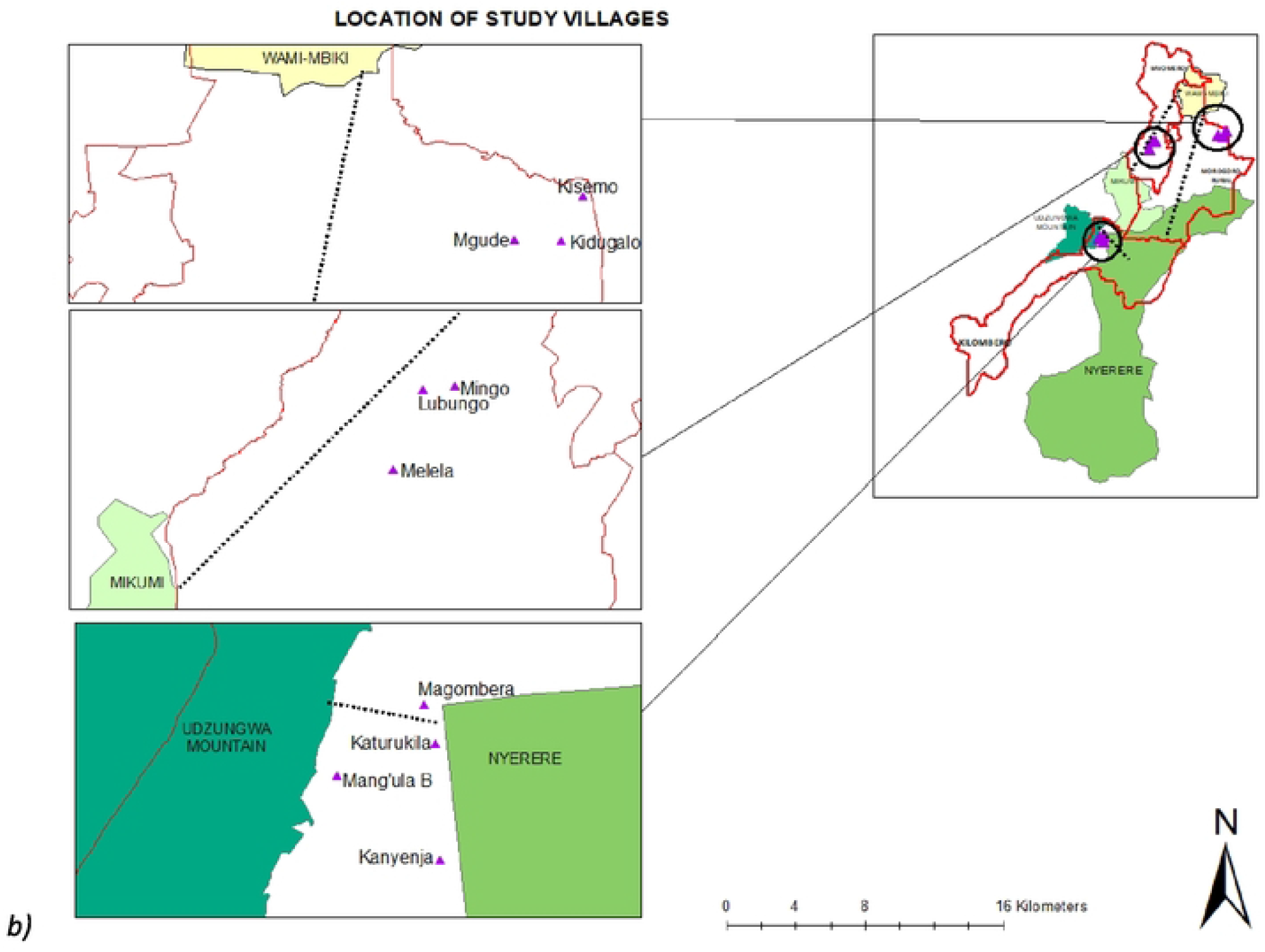
The location of the study area: a) the three districts of Kilombero, Mvomero and Morogoro Rural, and the nearby protected areas; b) The study villages in each of the districts.

Two of the three districts feature corridors connecting the protected areas to Wami-Mbiki WMA (see Figure 3), an important link to four different protected areas in Tanzania (Nyerere and Mikumi National Park to the south and southwest, and Handeni Game Reserve and Saadani National Park to the north and northeast, respectively). As areas of natural habitat decline, such corridors linking protected areas and other occurrences of natural habitat are important components of broader landscapes that enable movement of species across larger areas (23,27,28).

The Morogoro Region experiences a subtropical climate with distinct rainy and dry seasons. Although all three districts examined contain numerous streams and tributaries, a large portion of the Kilombero District lies in the Kilombero River Basin, located on the upstream part of the Rufiji River Basin. The Kilombero River Basin is agriculturally rich and attracts investments in the valley, many of which utilize water from the river and smaller streams for irrigation. The predominant economic activity in the study area is agriculture, with the majority of households practicing farming (80%) and a few herding livestock (29). Farming involves small, medium, and large tracts of land and include both rainfed and irrigated crop production. However, there was a shift in recent years towards more medium-scale farming associated with use of agrochemicals, where rice, peas, potatoes, cassava, maize, beans, and banana are food and cash crops; and sesame, sugarcane, sunflower and cacao are strictly grown commercially (30). In the Kilombero District, for example, commercial agriculture has increased rapidly in recent decades, leading to high deforestation that threatens biodiversity (26). Large-scale agricultural investments include those associated with sugar (Illovo Sugar Company) and rice (Kilombero Plantations Limited) production. Moreover, privatization policy in the mid-1990’s has had negative impacts on the valley, pushing subsistence farmers into marginal lands (31). Large-scale investments on land affect local people’s livelihoods through reduction in the amount of land available for subsistence agriculture and causing settlement displacement (26). The recent national agricultural strategy called the Southern Agricultural Growth Corridor of Tanzania, recipient of considerable international investment (32), is increasing large-scale crop production in this part of Tanzania. Local people are being gradually pushed to the outskirts of the valley, often closer to nearby protected areas where they experience increased HWC.

### Interviews and Data Analysis

To understand attitudes and determinants of changing relationship between people and elephants, we conducted surveys of 10 villages in three districts to gather information about their experiences and attitudes towards elephants (see S1 and S6). We used the conceptual model shown in Figure 1 to structure and design survey questions. The study examined villages in Kilombero, Mvomero and Morogoro Rural districts with the highest reported incidents of HEC for 2019-2020:

- Mvomero District: Melela, Mingo and Lubungo villages.
- Kilombero District: Kanyenja, Katurukila, Mang’ula B and Magombera.
- Morogoro Rural District: Kisemo, Mgude and Kidugalo.

For each of the above communities, we obtained village profile information for total population, number and names of sub-villages, main economic activities in each village and main livelihood challenges. In each village, the study surveyed a random sample of households. Based on inquiries conducted in similar settings (33)we used a minimal sample size of 5% of the total households, or 30 households per village in cases where 5% yielded fewer than 30 households. In all, the project interviewed a total of 454 households. Since the study involved human subjects, we obtained Institutional Review Board (IRB) clearance from the Pennsylvania State University (see S2).

We also conducted in-depth, semi-structured interviews of a subset of 180 villagers in total for the entire study area. In addition, we conducted semi-structured interviews with a total of 12 experts, including agricultural officers, game officers, land planning officers and community development officers from each of the three districts, purposively selected to include a range of experts in all sectors relating to the subject matter of this project (S3). The guiding questions for the semi-structured interviews appear in supporting documents (S6). We used Open Data Kit (ODK), a data collection tool that uses mobile devices and enables data sharing through an online server, to compile data from the study area.

We calculated descriptive statistics and conducted statistical analysis of the surveyed data using Statistical Package for the Social Sciences (SPSS). The survey included 41 variables that could have been included in statistical analysis of attitudes towards elephants, drawn from the questionnaire. We employed Principal Component Analysis to determine the factors with stronger contribution towards the variation in results. This reduced the variables to 13, all of which were included in forward stepwise binary logistic regression analysis as explanatory variables. We used Hosmer and Lemeshow Test to evaluate the model fitness and identify the variables influencing the variation in the dependent variable. The attitudinal response variable (dependent variable) was derived from a 5-point Likert scale response to the question “In a scale of 1-5 (1=I like elephants very much and 5=I hate elephants), what is your feeling towards elephants?” Given that there were very few responses in “like” category, we collapsed the 5-point scale into a binary variable, with categories reflecting an overall positive attitude, for instance “I like elephants very much,” “I like elephants” and “I moderately like elephants” grouped together as positive and the remaining answers (“I dislike elephants” and “I hate elephants”) categorized as negative. We used stepwise binary logistic regression to characterize the relative importance of all 13 explanatory variables in shaping attitude towards elephants among local communities in the study area.

The survey also assessed the level of tolerance of local communities towards elephants using the same set of variables as above. We based the dependent variable on the question “Which statement best describes your tolerance towards elephants: i) I tolerate elephants in my environment ii) I would tolerate elephants in my environment if they stopped destroying crops or iii) I would prefer elephants to be eradicated. We categorized these answers as “Tolerate,” “Conditionally tolerate” and “Eradicate” to indicate different levels of human-elephant coexistence with elephants in the study area. To determine what variables, influence respondent tolerance towards elephants, we used multinomial logistic regression with likelihood ratio test, a robust approach where the dependent variable has more than two categorical responses.

## Results

Survey results address HEI in villages that all face like challenges. All 10 communities have similar socioeconomic characteristics. Most residents practice subsistence agriculture with no alternative livelihood. This pattern is reflected in their monthly earnings, where most households earn less than US$100 per month. Likewise, most of the respondents in all surveyed villages had only primary education or no formal education. Experiences interacting with elephants also had similar patterns in the 10 villages. We present results largely in tabular form, with much of our examination of those results found in the discussion section below.

### Change in the attitude towards elephants

Over the past decade, there has been a shift in the local resident attitude towards elephants, primarily changing from favorable to unfavorable. The major change (81.9%) was from positive to negative, with 2.2% changing from strong positive to fairly positive (Table 1). In contrast, 15.9% of the respondents did not change their attitude, 9.5% remaining positive and 6.4% negative. Out of 382 whose feelings have changed, 303 said their feelings changed during the past five years, while 69 said their feelings changed between six to 10 years ago, and the remaining 10 said their feelings changed over 10 years ago.

**Table 1:** The change in feelings towards elephants expressed by the respondents.

|  | CHANGED ATTITUDE |  | NOT CHANGED ATTITUDE |  | TOTAL |
| --- | --- | --- | --- | --- | --- |
|  | From positive to negative | From strong positive to less positive | Remained positive | Remained negative |  |
| **Numbers** | 372 | 10 | 43 | 29 | 454 |
| **Percent** | 81.90 | 2.20 | 9.50 | 6.40 | 100 |

The binary logistic regression identified seven variables affecting the variation in attitudes. Six of those variables were statistically significant in explaining the variation in people’s attitudes towards elephants (p=0.05; Table 2). The model fit the data very well (p=0.002). The seven explanatory variables accounted for 78.9% of variation in the dependent variable.

**Table 2:** Variables Influencing Variation in Attitude in Kilombero, Mvomero and Morogoro Rural Districts.

| Variable | Explanation |
| --- | --- |
| *Whether feelings towards elephants have changed over time* | Individuals whose feelings have changed over time are more likely to have negative attitude toward elephants) compared to those whose feelings have remained the same (p = 0.000). |
| *Amount of crop lost to elephants* | Those who lost a half or more of their crops to elephants last year are more likely to have negative attitude towards elephants than those who lost one third of their crops to elephants (p = 0.000). |
| *Amount of crop lost due to other reasons* | Those who did not lose crops to other reasons were more likely to have a positive attitude towards elephants than those who lost one third of their crops to other reasons (p = 0.027). |
| *Personal readiness to tolerate elephants* | Individuals who said they would tolerate elephants in their environment if the elephants stopped destroying their crops were more likely to have negative attitude towards elephants than those who said they tolerate elephants in their environment even with the destruction they cause. (p = 0.000). |
| *Benefits from elephants* | Those who said did not receive any benefits from elephants are more likely to have negative attitude toward elephants than those who said they received benefits from elephants (p = 0.023). |
| *HEC Trend in the past 30 years* | Those who believed that HEC increased during the past 30 years were more likely to have negative attitude toward elephants than those who said they did not know (p = 0.0295). |

We did not find any statistical significance in variation in the overall attitude towards elephants by gender, age, education, income, cropping cycles, or farm size. When we examined these same data for individual districts, the model did not return any significant variables for Kilombero and Mvomero districts. Two variables emerged as statistically significant in influencing the attitudes in Morogoro Rural District: *Whether feelings towards elephants have changed over time* and *benefits from elephants.*

### Tolerance Towards Elephants

The multinomial logistic regression used to evaluate tolerance towards elephants using the variables above fit the data well (Likelihood Ratio Test = 73.3 %, p<0.001). As with attitudes, evaluation on the level of tolerance towards elephants showed no difference by age, gender, cropping cycle or participation in elephant conservation activities. However, there was a difference between those who earned less than 100 US Dollar per month, as they were more likely to have a conditional tolerance or prefer eradication of elephants than those who earned more (p=0.007 and p=0.009). In addition, those who believe that the community is not living peacefully due to escalating HEC were more likely to have a conditional tolerance or want elephants eradicated (p=0.003 and p=007). Although not statistically significant, respondents who did not lose any of their crops to elephants were more likely to have unconditional tolerance towards elephants. Those who did not have any formal school education were more likely to tolerate elephants unconditionally (p=0.001) (Table 3).

**Table 3:** Multinomial logistic regression results for local communities tolerance towards elephants.

| Variable | B (S.E) | Wald | Exp(B) (95% C.I) | p-value |
| --- | --- | --- | --- | --- |
| **Conditional Tolerance** |  |  |  |  |
| Crops lost to elephants (*none*) | -11.042 | 5.926 | 1.601E-5(2.204E-9 – 0.116) | 0.15 |
| Education (*no formal education*) | -16.000 | 115.676 | 1.126E-7(6.098E-9 – 2.078E-6) | 0.000 |
| Perception on how community coexists with elephants (*unpeacefully due to escalating HEC*) | 4.122 | 8.699 | 61.670(3.986 – 954.113) | 0.003 |
| Income (< \$ 100) | 11.744 | 7.249 | 125986.407(24.402-650457428.6) | 0.007 |
| Time of the day (*anytime*) | 1.158 | 0.389 | 3.184(0.267- 37.920) | 0.360 |
| **Eradicate** |  |  |  |  |
| Crops lost to elephants (*none*) | -11.690 | 6.502 | 8.374E-6(1.048E-9 – 0.67) | 0.11 |
| Education (*no formal education*) | -14.387 | 112.733 | 5.648E-7(3.967E-8 – 8.040E-6) | 0.000 |
| Perception on how community coexists with elephants<br>(*unpeacefully due to escalating HEC*) | 3.643 | 7.360 | 38.188(2.748-530.612) | 0.007 |
| Income (< \$ 100) | 10.632 | 6.818 | 41434.403(14.175-121111090.2) | 0.009 |
| Time of the day (*anytime*) | 1.690 | 0.697 | 0.373(0.037-3.776) | 0.404 |

“Tolerate” was set as the reference category. Deviation from the “tolerate” is shown in the table above. The significant variables are shown in gray shades ( $p=0.05$ ). B (S.E) =Standard Deviation to the mean; Wald=Individual predictor; C.I=Confidence Interval.

## Discussion

Over the past decade, the attitudes of the vast majority of people included in this study have changed from positive to negative, primarily following escalating cases of crop damage and threats to human life. Based on the results, the communities perceive that HEC has become more severe in the past five to seven years. Overall, 88.3% of the respondents expressed a negative attitude towards elephants. A follow up question on whether feelings have changed or remained the same overtime revealed that most people whose feelings towards elephants changed went from positive to negative, the main reasons for the change being crop raiding and concerns for personal safety (see Table 1). Willingness to tolerate elephants is being undermined by increasing crop raiding. More negative interactions with elephants increases vulnerability in local community livelihood, potentially increasing the threat to elephants due to possibility of retaliatory actions. Attitudes and behaviors are linked, and while they may not be synonymous, they have the potential to influence each other (34). Inquiries into acceptance and tolerance of wildlife indicate that both phenomena are characterized by inaction of individuals when first affected by wildlife, where people would tolerate wildlife until it reaches a point where individual, or society inaction ceases, and behavior intended to affect wildlife negatively begins. When situations reach this threshold, intolerance begins (34).

Human behaviour in interacting with elephants can thus be arrayed along a continuum from acceptance and tolerance of elephants, possibly taking the form of stewardship or inaction, to intolerance that can take the form of actions intended to harm elephants. This study finds evidence of such a shift in villages from the three districts examined. The confidence of villagers to express vocally their unfavorable attitude towards elephants indicates that a threshold towards intolerance has been crossed.

Based on the statistical analysis, discussion with the local communities and experts, as well as literature review, a number of factors emerge as important social-ecological determinants of the nature of human-elephant relationship (Figure 4). These factors belong both to the ecological and social systems involved and relate to tolerance-attitudes towards elephants and to the intensity of conflicts between people and elephants. We examine results as they pertain to these two systems.

**Figure 4:**
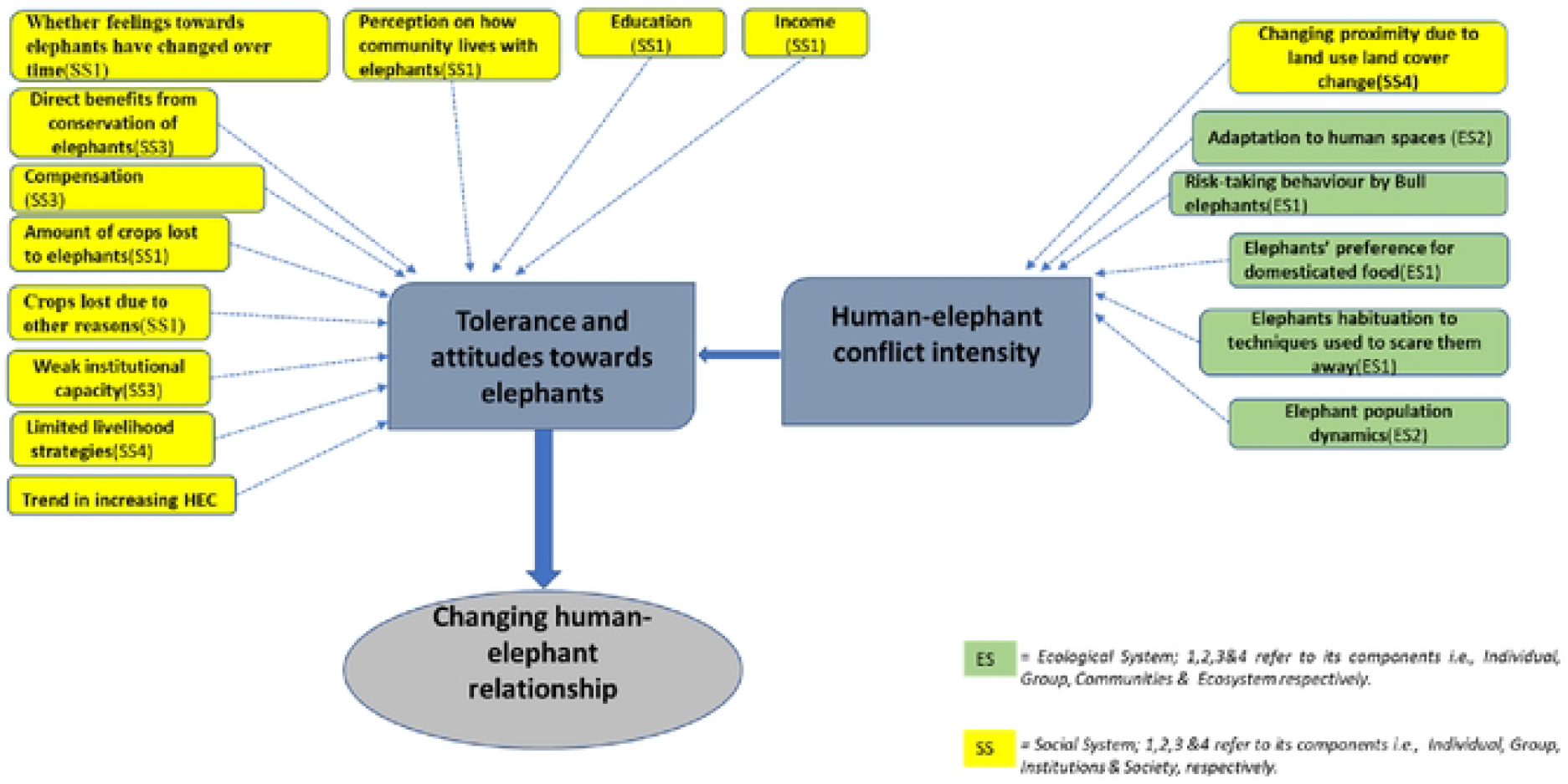
Social-ecological determinants of human-elephant relationship. SS=Social System and ES≈Ecological System. SS and ES and their associated numbers correspond to SS and ES components in the Social Ecological System framework in figure I above.

### Social Aspects

#### i. Trend in Increasing HEC

Not surprisingly, the community members who believe HEC increased during the past 30 years were more likely to have negative attitudes toward elephants. These are people at least 30 years old who have lived for that long in the village. The community members attributed the increase in HEC to growing human population that now lives in previously uninhabited areas causing more frequent encounters with elephants. Parts of some villages are reported to occupy wildlife corridors between protected areas (see Figure 3), although the villagers contested this idea, completely denying existence of any corridors.

*“We were compensated every 1 hectare for 75, 000 TZS* [Tanzanian shillings] *because we were told we cultivated in the reserved area. We were told to move out of the corridor. We didn’t understand how the village became a corridor area,”* complained a woman in Mgude village. *“There are no elephant corridors in this area, in the past 50 years there weren’t any corridors, we have been surprised by the sudden presence of elephants*,” added another woman.

The discussion with the District Game Officers (DGOs) revealed that villager denial of corridor existence is linked to the fear of losing land, since community members have seen how people who encroached protected areas in other places were forcefully removed. One main problem is that Tanzanian wildlife authorities did not gazette or formally mark and protect wildlife corridors, leading to the potential for land use conflicts with the villages along such corridors.

Villagers report that HEC has increased in the past five or six years. *“I am now 77 years and all my life I haven’t seen what elephants are doing now, in the past they didn’t do this…these elephants have come about six or five years ago. In the past we were dealing with wild pigs only, but now we have elephants, more and more elephants coming to the village,”* a woman in Mgude village reported. *“HEC has increased in the past five years, it looks like elephants’ number has increased. In the past, when I was a kid, we didn’t have a problem with elephants. We didn’t see them as often, but now we see them every day,”* reported a man in Kidugalo village. Another man added “*Human population has increased. We have encroached into wildlife spaces (corridors), so elephants have no option ….”*

In line with the local communities’ observation, Tanzania Wildlife Research Institute (TAWIRI) reports revealed that recently elephant population in the Selous-Mikumi ecosystem has slightly increased(35). Conservationists in the area attribute the increase to decline in elephant poaching in recent years; this possibly has led to elephants being confident to enter villages and village land without fear of being killed. Another possibility is that, with climate change and environmental degradation, elephants are forced to move between habitats more often than before, leading to more frequent encounters with people, particularly when elephants leave protected areas. Confirming this claim would require a systematic study of elephant habitat inside the protected areas, which is beyond the scope of the current study, although it could help to explain why elephants are venturing into villages when they did not do so in the past.

#### ii. Lack of Compensation

In Tanzania, the Wildlife Conservation Act No. 5 of 2009 and its recently reviewed 2018 policy does not provide for compensation to local communities whose crops are damaged by wildlife. Although there is a provision allowing people to kill wildlife for self-defense or in defense of their property, this only applies under hunting activities and pastoralists in defense of their livestock; it does not extend to allow farmers to kill crop-damaging wildlife (20). Under the Ministry of Natural Resources and Tourism, wildlife found outside the national park boundaries are under the care of Game Department. Under certain situations, officers of the Game Department are allowed to kill wildlife that is causing problems within human spaces. The Wildlife Act, however, provides for “consolation” payment to farmers in case of crop raiding or injuries and loss of life caused by wildlife. Consolation amount does not cover the loss incurred but serves as a gesture to sympathize with the victims of wildlife damages. How does the consolation work? The experience of farmers in the study area and elsewhere in the country tell a story of how complicated and bureaucratic process it is to obtain the consolation payment. According to the DGOs, the amount provided for one acre of crops damaged by wildlife is 25,000 Tanzanian Shillings (~US$11), regardless of the type of crops, or production cost incurred. For loss of human life, consolation amount is 1,000,000 Tanzanian Shillings (~US$440) to the victim’s family. This amount has been highly contested, leading to a perception that wildlife is valued more than human life, considering that the fine imposed on elephant-poaching criminals for one poached elephant is US$15,000 - US$30,000. *“In this country elephants are valued more than people, if elephants kill you or a member of your family, the maximum you can get, after a long process is just one million shillings, but if you happen to kill an elephant, you will be fined millions of money, and you will face jail.”*

For subsistence agricultural communities whose livelihoods directly depend on what they grow, and whose access to natural resources is restricted, different values placed on wildlife and humans have a direct bearing on economic and psychological costs of living with wildlife. Given that villagers have invested significant time and effort in livelihood strategies that are now threatened by wildlife, the authorities not placing equal value in compensation for wildlife-caused damage likely will maintain negative attitudes toward wildlife (36)). *“We wanted to form a special committee of elders to go and see the president to tell him about our problem…there is no food…and no compensation…just last week a person was killed by an elephant…about 16 people (in this ward) have already been killed by elephants,”* complained one elder from Mingo village. *“I was born in 1940 … there were no elephants in this village … but now elephants are rampant … last year I planted two acres but didn’t harvest anything … when Nyerere was president, when Mwinyi was president, Mkapa even until Kikwete the situation was not that bad … but now the situation is terrible.what will I do as an elder… they have promised compensation but we have received nothing”* an elder of Melela village remarked.

According to the Morogoro Rural DGO, the last compensation made in that district was in May 2021, for those who lost their crops to elephants in 2019. The government has paid a total of 56.3 million TZS (US$24,415) as consolation in Kisaki, Ngerengere, Mgude, Kisemo for crops lost in the district that year. To this, the agricultural officer commented that *“Yes, it is provided but it takes a very long time, the consolation might take even two years to be made, and always … out of the season, by the time it comes, the person has suffered a lot, it does not reflect the loss incurred. Farmers are left frustrated, not knowing what to do for the season the loss is incurred … Imagine if somebody was injured and does not have money to go to the hospital? They may die while waiting for consolation money,”*

#### iii. Weak Institutional Capacity

Game officers’ capacity to respond to crop raiding incidents is very limited. The Kilombero DGO offered, *“Employees of the Wildlife Section are very few. I cannot respond alone when I hear a group of poachers or some people are in possession of bushmeat, or even elephants invading people’s fields, I have to mobilize resources first before I go. This delays the response. If you call the police department most of the time, they also do not have transport … you may ask for a motorcycle from someone but still you have to get fuel. I cannot blame the district executive officer, because they are also allocated restrictive budget which does not give priority to conservation issues.”* A similar message was echoed by the Mvomero DGO: *“…The situation will get worse … it is undermining my confidence and they* [local people] *are becoming hostile against us* [conservationists]*, and maybe in the future they will start stoning us … even when I go to the villages, they always shout that I should leave with my elephants. The ministry of natural resources lacks adequate capacity and resources to respond to human-wildlife conflicts.”* The DGO in Morogoro Rural District shared similar experience that the department’s capacity is very low to respond to incidences of HEC “*We have very few human resources, for example, here at headquarters I am just alone. Also, transport is a challenge, when I am called to respond to an incident, I fail to respond timely because I do not have a motor vehicle that is on standby. Even our collaborators, TAWA and TANAPA* [Tanzania National Parks Authority], *when I call them to help, they always say their patrol groups are responding to a different district, and they do not have extra human resources nor transport to provide us. Because they are responsible for the entire Morogoro Region, responding also to Kilosa, Mvomero …”*

Villagers similarly made comments about limited institutional capacity. In Kanyenja, villagers reported, “*TANAPA do not have anything, they do not have equipment. Sometimes they come here and run away without doing anything.”* In Lubungo the villagers complained *“Game people are overwhelmed, we call her here to chase away elephants, before she does anything she is already called in Mingo village … elephants are raiding there also … she needs more people to work with.”*

Inability to respond timely to alerts from farmers about elephants invading their fields potentially increases tension between people and elephants. Considering the special protection status of elephants, coupled with their huge body size and frequently aggressive behavior, a farmer cannot confront them and is left with no option but to wait for action from responsible authorities. When the desire to save their crops from elephants is not met due to delays in response, tolerance is undermined further and contributes to the difficulties of coexistence for these two species. Such problems extend to local (village) leaders, as stated by the Melela village chairperson “*I have planted tomatoes … two acres. The elephants have eaten them all … when I was actually there looking … but I couldn’t do anything. We do not have any means to protect ourselves from elephants.” “We call the game people, when they come and start dealing with them, and more elephants come, they run away, am I lying?* In response, all members replied *“no, you are not lying.” “They do not have the capacity to control them. The only way is to harvest them…,”* complained a woman in Mgude village. In the Kilombero District, the DGO’s remarks confirmed this shortage *“UMNP* [Udzungwa Mountains National Park] *at times provide[s] the DGO with their vehicle, but they cannot always depend on them as they also have their own plans … The human personnel in the wildlife department are very few, I cannot respond for example to a group of poachers by myself”.*

#### iv. Crops Lost to Elephants

Local community readiness to share space with elephants is degraded by damage from elephants, especially in agricultural fields. Our household surveys showed that 63.9% of the project sample lost at least half their crops to elephants during the previous year. These people understandably have negative attitudes toward elephants. The resentment about crop raiding reflects the struggle of the local communities to sustain livelihoods through subsistence farming. In all the surveyed villages, elephants were reported as the major challenge to agricultural practices, along with disease and pests. Crop damage by elephants compounds existing livelihood challenges and is a major impediment of poverty alleviation. *“We plant big farms, but we don’t harvest anything, the elephants come and harvest everything, leaving us hungry, this is the biggest challenge we have,”* complained a woman in Mgude village. *“We have no income due to not getting enough harvest since elephants eat our crops … livestock also eat our crops. We are left with nothing for food, nor fees nor money for medical care,”* added another resident of that village. *“How can we take our kids to school? Tomatoes, sugarcane, cassava, are the ones giving us school fees … but we are not harvesting,”* complained a woman in Kidugalo village. Similar sentiments were expressed in Kisemo village: *“We are forced to harvest before the crops are fully ready, because we want to prevent them from being damaged…but this causes the produce to go bad.”* Crop damage by elephants leaves subsistence farmers despondent and resentful towards elephants. Owing to their gigantic sizes and big appetites, elephants can destroy a family’s crop overnight, leaving them with no food for the coming year. Many community members, particularly women, borrow money from a local village community bank to invest in farming— to purchase farm implements and inputs, as well as agrochemicals to fight diseases and pests, hoping to pay back the loan after harvesting. Usually, they would sell part of the harvest to pay back the loan and use the rest for food and the upkeep of the family. After villagers invest their resources and effort in farming, elephants damaging crops in the fields leaves the households despairing and less likely to have favorable attitude towards elephants. If the current situation continues, the farmers fear they will fall back into terrible poverty and be unable to care for their families. *“Farmers make investment with no return, they are forced to go and borrow money from financial institutions for their farming, and if the crops are all damaged, they may find themselves in a terrible situation and are left with debts, making them fall into abject poverty,”* said the agricultural officer in Morogoro Rural District. In Katurukila some elders added, *“We are only sure of planting, but not harvesting.”*

#### v. Crops Lost to Other Reasons

Those who have lost crops to elephants and to other reasons were particularly bitter about elephants. The results suggest an existence of some threshold, with loss of one third of the crops to other reasons triggering negative attitude towards elephants. Given that the survival of subsistence farmers depends on what they harvest from their farms, losing a third of crops to other reasons and adding elephant crop damage on top of that, affect them significantly. “Other reasons” in this case primarily refer to diseases, crops failure due to bad weather, and rodents eating stored harvest. However, it seems that the blame sometimes is attributed to the visible and bigger cause of the loss, in this case, elephants. To some extent, this makes sense since more loss of crops compounds the situation. Instances where some crops were lost to diseases and pests at an earlier stage, and surviving crops matured only to have elephants destroy them just before harvest, amplifies farmer frustration and disappointment and, ultimately, resentment towards elephants.

#### vi. Benefits from Elephants

Benefits from elephants have the potential to influence human attitudes towards elephants. Individuals in the communities recognize, realize, and interpret both tangible and intangible benefits from elephants differently. Most people do not perceive elephants as having a direct benefit to them. Although they acknowledge that elephants have intrinsic, aesthetic and touristic value that benefit the country, they do not believe these benefits affect them locally and feel they come at the cost of HEC which the local communities suffer. *“Elephants are important for tourism, but they should be prevented from coming to our villages by putting a fence around the villages near protected areas…They are telling us not to go in the protected areas, but they should also play their part by preventing elephants from causing damages. The government is getting benefits from elephants, but we get a loss,”* complained a woman in Kidugalo village.

The presence of elephants and the revenues from tourism to see them and other attractions has brought some tangible, collective benefits to local villages such as building school classrooms and dispensaries, purchasing school desks and contributing to social funds by Mikumi, Nyerere and Udzungwa Mountains national parks. However, the amounts of crops lost to elephants affects household’s food security and outweighs the collective benefits of such gestures. Social funds and consolation money for crop damage do not offset the costs incurred from HEC. A more complete cost-benefit analysis would provide the basis to make tangible benefits from conservation adequate to improve local attitudes towards elephants.

#### vii. Personal Tolerance

According to the DGOs, the communities still show some cooperation when it comes to conservation activities, but it may not be long until they stop because their tolerance is disappearing. The analysis presented above suggests that for many villagers tolerating elephants in shared landscapes would require reduced crop destruction. These individuals were more likely to have negative attitude towards elephants than those who said they tolerate elephants in their environment even with the destruction they cause. The latter group consisted mainly of the few households that had an alternative livelihood strategy, such as a small business. HEC may trigger economic, social and psychological impacts on local communities experiencing it, such as food insecurity, poor sleep, increased exposure to malaria and dangerous animals at night while guarding fields and poor school attendance (36),(37)). Unfortunately, tolerance towards elephants likely will erode further if the current situation continues, which will have negative implications for rural villages and for biodiversity conservation (and the tourism associated with it). Experience shows that where tension is high and tolerance thresholds surpassed, elephants suffer retaliatory killings from local communities, in some cases even leading to collaboration with poachers in killing elephants (38) (39)

#### viii. Limited Livelihood Strategies

Reliance of communities in the study area on subsistence agriculture as their primary livelihood is central to the survival of most households. The majority surveyed earned less than US$100 monthly, and less than 3% had an alternative livelihood strategy. Over-reliance on a single source of livelihood increases household vulnerability to external shocks. For farming communities, resilience of such livelihoods is substantially undermined in the face of weather extremes, diseases and pests or crop raiding. In contrast, communities which have a backup source of livelihoods are less affected by crop damage by wildlife and have a more favorable attitude towards wildlife (see Table 3). Situated near protected areas, local communities in this study are restricted from accessing timber and non-timber forest products such as firewood, charcoal, fruits, or honey from the reserves, which could otherwise contribute to generating alternative income. Lack of livelihood diversification can potentially lead to maladaptive responses such as trying to increase crop yields through expanding croplands into elephant habitat that will amplify HEC. Beyond near-term risks, this will further undermine livelihood sustainability, reducing the ability to cope with and recover from shocks and enhance its capabilities without compromising the natural resource base (40)

#### ix. Education

Surprisingly, individuals with no formal or only primary education were more likely to tolerate elephants unconditionally compared to those with a higher education level. There are two possible explanations for this. First, people with more formal education likely have a better understanding of the amount of money elephants bring to the country through tourism and may be sending a message that such financial benefits come at a cost that they are unwilling to cover without greater compensation. Second, the younger generation, who are more likely to have acquired secondary and college education are more likely to be persuaded by politicians who often drive the agenda against conservation to secure votes. Discussion with ecologists in Nyerere National Park, UMNP and Mikumi National Park highlighted that HEC is now being used as a political agenda, “*We are now receiving calls from the members of parliament or madiwani* [ward representatives]*, telling us to go drive elephants from the villages. We understand the problem is there, but the politicians are not helping, they are fueling the situation, and raise false hopes and expectations. They are greatly undermining people’s tolerance. Just last year the minister* [of Natural Resources and Tourism] *toured the whole country, telling people ‘We have now found a panacea for HEC, the government will be providing financial compensation,’ something which is not practical. The government cannot compensate for all elephant-caused damage because it’s too expensive and complicated.”* remarked one of ecologists. The youth and the younger generation can easily follow, and access rhetoric provided by politicians through social media platforms. They can easily be recruited against conservation of elephants because many of them are jobless. The high unemployment rate in Tanzania, which forces graduates to return to their villages, complicates matters, making youth bitter and intolerant against any crop loss to elephants. It renders them prepared to believe politicians supporting certain positions and easy targets in driving a political agenda.

### 2.4.2 Ecological Aspects

Exploring and understanding behavioral, physical, demographic or personality traits of elephants is critical in understanding the interaction between people and elephants. Biological and ecological research provides increased understanding of elephant behavior to help identify the ecological factors determining human-elephant relationships in shared landscapes. Given the challenge of conducting long-term field observation of elephants, I complemented the data obtained from surveys, focus group discussions and expert interviews with literature review on elephants and their interaction with humans from studies conducted in the case study area as well as in other relevant areas. The following issues emerged as important ecological topics influencing human-elephants’ interactions in the study area.

#### i. Elephant Population

Elephant population in the Selous-Mikumi ecosystem declined dramatically following years of serious poaching (20).The southern Tanzania elephant population is relatively young and has been subjected to heavy poaching beginning in the 1970’s and 1980’s. Between 1976 and 2013 elephant population in the Selous-Mikumi ecosystem declined from 109,000 to 13,000 (41). In some habitats, such as in UMNP, poaching threatened extinction (20). A TAWIRI report indicates that the Selous-Mikumi ecosystem elephant population has started stabilizing, with a marginal increase from 15,217 in 2014 to 15,501 in 2018 (20). The gazettement of the UMNP allowed some recovery of elephant population in the area. However, even modest increases elephant population, the totals well below historic levels, still present challenges in the form of potential conflicts with local communities. The Kilombero DGO, who has worked in the district since 2011, confirmed that as elephant population in the Selous-Udzungwa-Mikumi has stabilized and slightly increased, HEC has increased: *“We must admit that to some extent elephants have increased partly due to declining poaching. Now HEC has increased. Crop raiding has increased.”*

#### ii. Individual and Group Behaviour

Individual elephant attributes also play an important role in HEC, contributing to changes in human-elephant relationship. Recent research (15)argues that an increase in HWC rates does not necessarily reflect growth in the wildlife’s population, since it could be a result of animals shifting their behaviour in response to changing environmental conditions. The following are some of the group and individual attributes associated with the behaviour of elephants that were considered to contribute to HEC.

##### a. Habituation to techniques used to limit their movements or scare them away

Elephants are intelligent social animals with high cognitive capacity. In the study area, where fences and physical barriers are constructed, elephants have been able to break through such obstacles and damage crops. One study (21) for example, explores the effectiveness of beehive fences in the eastern part of UMNP as a mitigation measure against crop raiding. Their findings show mixed results; in certain instances, beehive fences did not stop HEC completely as elephants destroyed parts of the fence. Also, presence of a fence only transferred the problem to nearby farms that are not so protected. Indeed, in Kilombero and Morogoro Rural districts where beehive fences exist in some localities, many villagers believe that such fences do not address the problem of HEC: *“Pepper, beehives, did not help, they just walk by and go to the next field, in fenced parts they break the fence and break the beehives before even the bees occupy them,”* said an elderly man in Kisemo village.

One important consideration is that elephants often become habituated to the techniques used to drive them away. They have become more confident roaming around and are not scared of many mitigation methods. *“Even when you beat the iron sheets they do not leave, in the past when we did that they left immediately, but now it is like you are playing a drum for them to dance, they won’t leave,”* said a woman in Kisemo village. *“We make noise and beat drums, flutes or trumpets and mix pepper with their dung etc. but they are used to all these, the methods do not help. Nowadays they are used to them, they don’t go away. They are not afraid of anything. …We don’t know what we should do anymore,”* reported a man in Kidugalo. A woman in Mgude village expressed similar sentiments: *“We use pepper, beehives, but after elephants realized that these don’t kill them, they started getting used to them. We have failed with all the methods; elephants have become more aggressive.”*

Even the use of bullets by game officers did not drive elephants very far, and after a few minutes observers reported that the animals returned and continued with crop raiding. *“In the past when we reported them to the authorities they were driven away and they would leave immediately, but now it’s like they are used to the bullets, neither are they scared, nor do they leave. They are literally camping in the village,*” said a man in Mingo village. The Kilombero DGO also confirmed this experience *“In the past we could chase them and wait for a few weeks before they show up again, but now they are habituated and don’t even go far, they return before you even leave the area.”*

The Morogoro Rural DGO also shared a similar observation about elephants’ ability to adapt to strategies designed to reduce HEC. “*Elephants have changed their behavior over time, they are becoming smarter I think … for example, for beehive fence, they avoid only the area that is fenced, if it’s 3 km they pass it and go to the next field. So, the fence is not addressing, but moves the problem to the next field.”*

It is evident that elephants are changing their behavior to adapt to existing mitigation methods and continue their crop raiding behaviour. As a result, this leaves the local communities frustrated, undermining their tolerance, and hence shifting their attitudes towards elephants to be more negative.

##### b. Preference of domesticated /grown food over their natural food

Elephants are generalist foragers with diverse diet including fruits, grasses, leaves, twigs, roots, bark and forbs (42). Within the study area, elephants have been reported to go for certain foods in human dominated space that are not available in the wild. Once they start eating crops as their food, they tend to prefer it over wild food, leading to more crop raiding (43). *“I was born in 1940 …there were no elephants in this village … but now even tourists contribute to their behaviour change as they give them food and they come for more in the villages*,” said a resident of Melela village. *“They are like the villagers, they don’t eat trees anymore, they eat cassava and fruits we have in our homes,”* Added another resident*.”* The DGO in Morogoro Rural District reiterated a similar view. *“They have changed their eating behavior: They are no longer interested in their wild food, now they are eating tomatoes, watermelon, pumpkin, rice, sugarcane, and cassava. Some crops they don’t eat but they trample over, for example sunflower, unripe papaya, unripe banana etc. I think they are angry when they don’t get what they wanted they angrily decide to trample over the rest that they do not eat or are not ready to eat.”*

The presence of agricultural landscape adjacent to protected areas significantly contributes to change in foraging behavior of herbivores in general (43). Increased preference for crops encourages crop raiding and hence influences negative attitudes towards elephants. A study in the Selous-Niassa elephant corridor found that crop damage by elephants was strongly influenced by the abundance of their preferred food crops, which included maize, bananas, potatoes, pumpkins, peanuts and onions (44).

##### c. Risk-taking Behaviour by Bulls

A recent study conducted along the eastern UMNP boundary between 2010 and 2014 used camera traps to identify elephants involved in crop raiding (22).Their findings show that adult males are more likely to crop raid and can pursue high risk feeding behaviour. They found many occasional crop-raiding elephants (32 out of 48) compared to repeat ones (16 out of 48). For male elephants, crop-foraging is a high risk, high-gain strategy to maximize the nutrients intake meanwhile minimizing foraging time and distance. Mumbi and Plotnik (45)link this behavior to fitness benefits since dominance and access to mates is related to body size.

Bull elephants leave their maternal families and form age mate groups who are more likely to engage in crop raiding for easy and fast nutrient gain while avoiding contact with older male elephants (46). Improved nutrition improves the chances that male elephants will have reproductive success (47)(48),(45) (49). Bulls are capable of developing crop raiding behavior through social learning; as a result, the tendency for raiding can be structured by male association networks, leading to different crop raiding behavior among individual bulls (46).

Prior to the study by Smit and colleagues (22), a similar study was conducted by Kabepele (20) to determine age, sex and identity of crop raiding elephants around UMNP. That research employed camera trapping, dung diameter measurement and genetics. Although the findings resembled those of Smit and colleagues (22), they also identified female elephants engaging in crop raiding. This was not captured on cameras, but through dung and genetic analysis. The findings confirmed that although male elephants dominate crop raiding, female elephants can also take risks in crop raiding though with high caution since they live in matriarchal family units with their infants and would therefore try to avoid risking the safety of their offspring.

##### d. Adaptation to human spaces

In all three districts, elephants have changed their behavior and are not afraid to maraud in villages even during the day. The DGO in Kilombero District reported that elephant behaviour has changed: *“In the past, about 7 years ago, elephants used to wait until nighttime, that’s when they would come out of the park, but now they are showing up during the day. In Msalise for example, at around 10 in the morning you may see them crossing the school compound while kids are in class. I think this is associated with changes in poaching. I think they are not charging and attacking people as much. They are not scared of people. But they could be triggered at any moment and attack people still.”* In Lubungo village, residents echoed similar sentiments: *“These animals have created a dwelling place here, day and night they are here, that is why we cannot work. When we tell the government we are told that we should get used to them, how can we get used to these animals? Why doesn’t the government get used to poachers if at all it is possible to get used to?”*

Increased presence of elephants in villages has become a great concern for residents. *“I was born in 1948 I didn’t see elephants in the village…until very recently, around 2018…it almost feels like they have all moved out of their areas and moved to this village…we cannot do anything to them, we have no capacity to fight them,”* an elderly man in Melela village said. Elephant confidence to roam in people’s residence and enter homes and marketplaces without any fear has increased, often bringing chaos. The increase in the number of confidently roaming elephants may reflect more restrictions imposed on resource access in Selous Game Reserve. Converting part of Selous Game Reserve to Nyerere National Park meant total prohibition of human activities in the area now designated national park. Reasons for repurposing part of Selous included controlling poaching, banning hunting and enhancing revenues from tourism, but on the other hand it brought sense of peace and freedom to elephants. *“Around 2007, it wasn’t that bad, but from 2015, 2017 elephants have increased in population so much. In the past when we were allowed to go to collect firewood from Selous, at least elephants would be scared of people and hide in the park, but since we were banned from accessing it, they have become freer. Now anytime they come and walk on the road like people, confidently,”* an elderly man in Mang’ula B village explained.

The presence of elephants poses security threats to school children, affecting their education, as reported in Katurukila village *“The schools are very far, and you can see elephant dung all over this way to school. Many times, our kids fail to go to school. They are young. Some parents have decided not to take their kids to school,”* complained a woman in Katurukila.

##### e. Elephants’ high olfactory capacity

Elephants have an excellent sense of smell, and when they smell fruits on the trees surrounding homes or stored inside the house, they often try to obtain them, often breaking into buildings. *“Last month they broke our neighbor’s shop. He had ripe mangoes in the shop, the elephants got the smell and destroyed the shop building,”* a woman in Magombera said. Having fruit trees around the house was also considered another risk for elephant attack, since elephants smell them from far and follow them. *“Just last week here in Magombera, an old man was killed at his house. He had mango trees around his house and there was no one at home when elephants showed up and started eating mangoes, so when he returned home and was trying to open the door, elephants came from behind his house and killed him,”* reported one Magombera villager.

#### iii. Changing Proximity due to Land Cover Modification

Over the past three decades land cover in the study area has significantly changed. Analysis of land use change between 1994 and 2020 indicate that human activities have expanded considerably, often replacing natural vegetation cover near protected areas as well as in elephant dispersal areas and migratory corridors. As a result of these land use/land cover changes, elephants and humans are coming into closer proximity, leading to an increase in negative interactions. Between 1994 and 2020, cropland expanded by 68% in the study area, where grassland, wetland and forests are being converted to agricultural fields (Figure 5). Also, some villages themselves have expanded to where elephants’ habitats are located. The DGOs and planning officials argued that cropland expansion is the main cause of increasing HEC, an opinion shared by many of the villagers interviewed in this study based on comments presented above.

**Figure 5:**
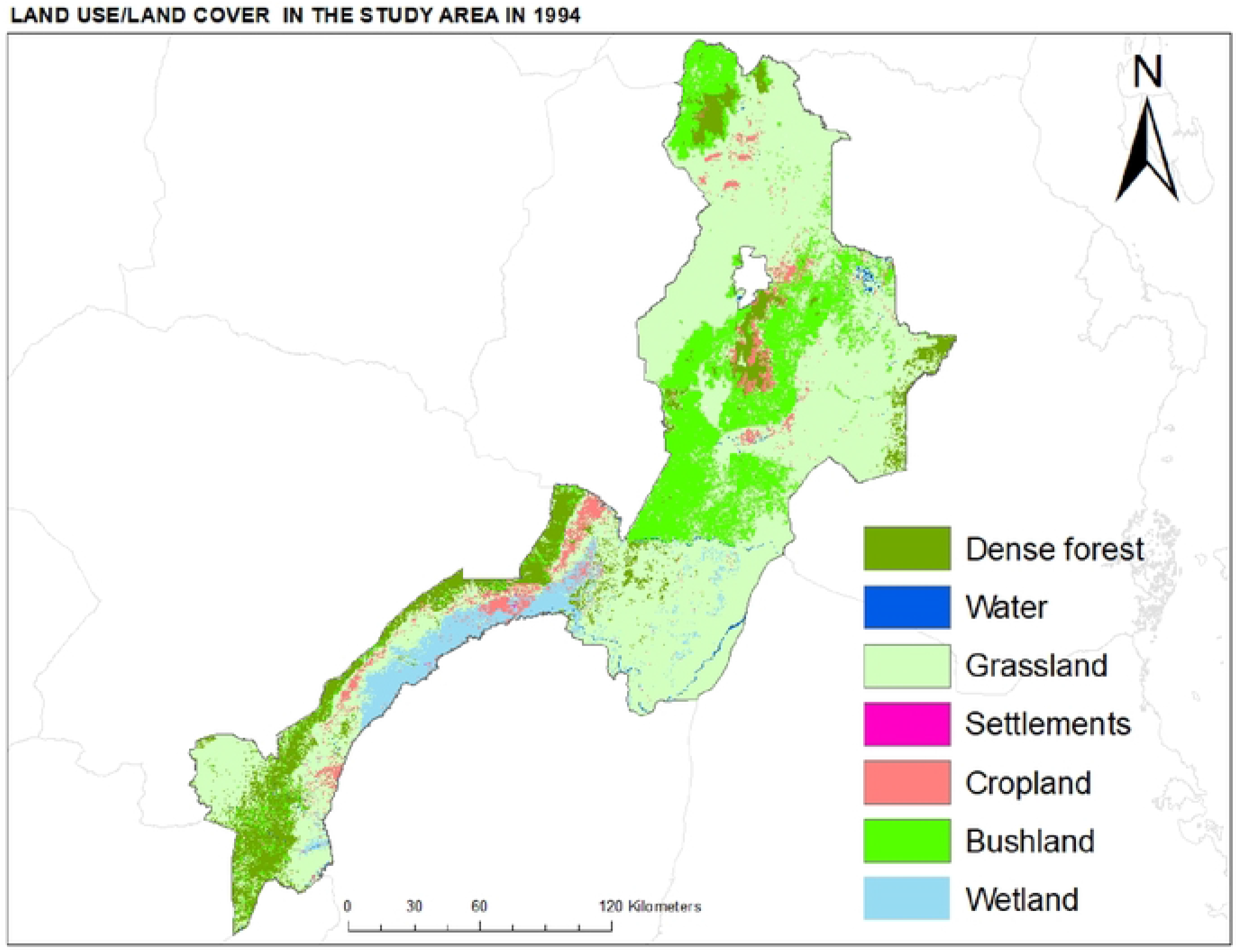
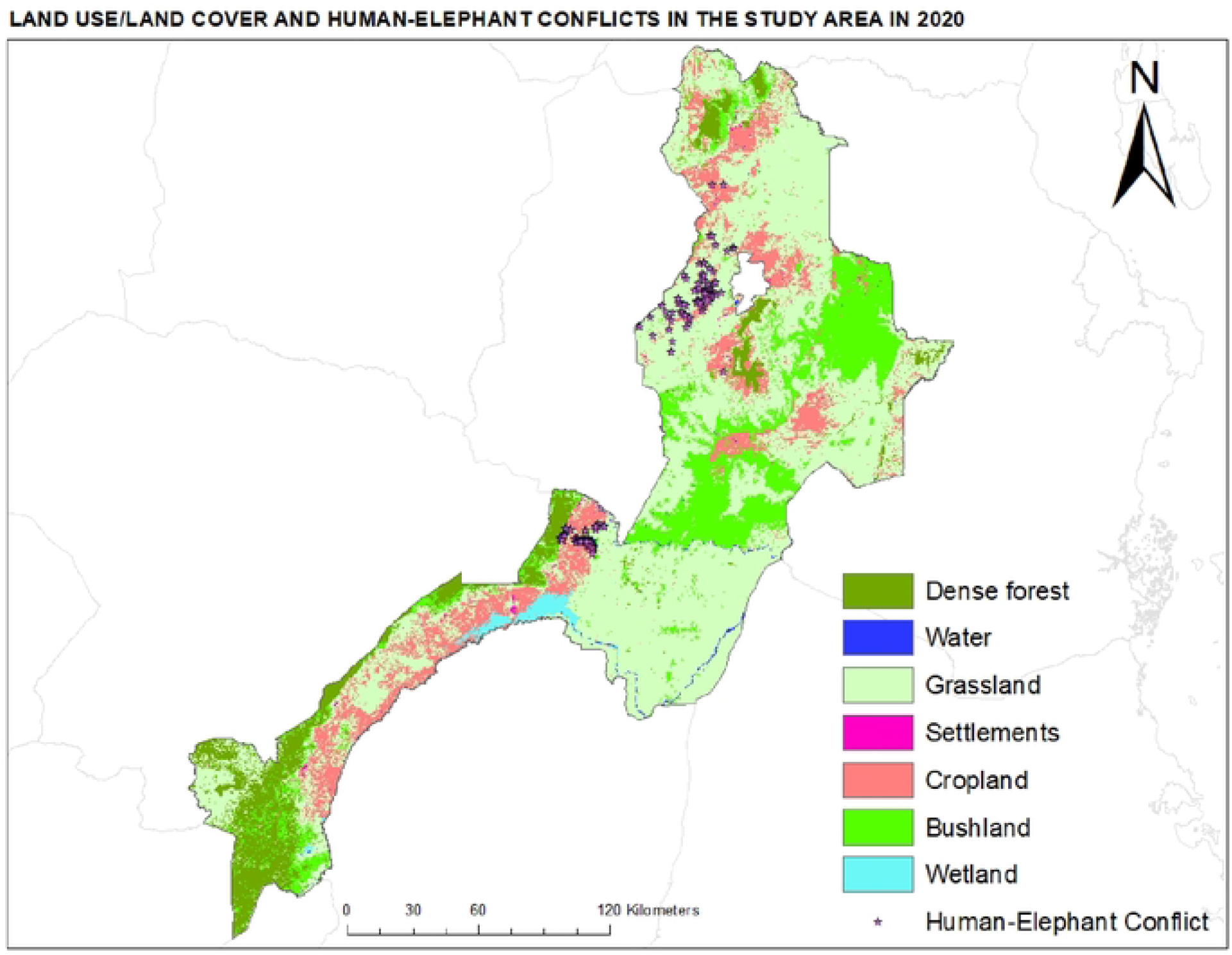
Land use/land cover mapping for the study area for 1994 and 2020 shown in seven distinct categories and locations for human-elephant conflicts in Kilomhero and Mvomero Districts.

### 2.4.3 CONCLUSION

Drawing from SES framework, our study of 10 villages in three districts in rural south-central Tanzania found a relatively recent shift in conflict-coexistence of humans and elephants from positive to negative, moving from minor to more severe impacts and from infrequent to more common interactions. An increase in HEC affects subsistence farmer tolerance, resulting in less favorable attitudes towards elephants. Social-ecological aspects help connect human perceptions and attitudes with ecosystems. On the social side, considerations shaping attitudes towards elephants include compensation, benefits accrued from elephants, amount of crops lost to elephants, amount of crops lost to other reasons, readiness to share space with elephants, whether feelings towards elephants have changed and changes in the social views of HEC in the past 30 years. On the ecological side, considerations affecting HEC and the human-elephant relationship in general include recent increases in elephant population, habituation of elephants to techniques used to scare them away, preference of agricultural produce over natural food, risk-taking behaviour by bulls, adaptation to human spaces, and elephants’ excellent sense of smell. In combination, these SES elements suggest continued conflict into the future.

Negative attitudes resulting from HEC present challenges for biodiversity conservation, development and livelihood improvement. HEC can potentially undermine other development strategies, for example through lack of response or support for collective development activities, fear in attending school and failure to conduct daily routines (16). Researchers argue that HWC in already vulnerable communities can exacerbate existing problems such as poverty, social inequality and feelings of oppression (36), (37).

Other studies have shown that the combination of these drivers can lead to retaliatory killing of elephants. In Kilimanjaro, Tanzania, for example, local people drove six elephants over a cliff to kill them, due to failure of district authorities to deter crop raiding by elephants, lack of benefits from elephants’ conservation, lack of compensation for the crops damaged, extra work incurred in guarding farms against elephants and protesting new elephant corridor imposition (38).

Human behavior in interacting with wildlife can be arrayed along a continuum from actions intended to harm elephants, to inaction to actions intended to benefit elephants, essentially moving from intolerance to acceptance/tolerance to stewardship. These shifts in attitude can be in either direction depending on existing situations. To improve tolerance level and to move up this continuum towards more positive attitudes that can transition into stewardship, we suggest increased intervention by the government and non-government actors to address some of the challenges faced by subsistence farmers in the area. This might take the form of subsidizing farm inputs and implements, improving the process of concession payments, or increasing the reaction of local government agencies to incidents of HEC. These will reduce the cost incurred in farming, possibly motivate subsistence farmers to tolerate crop loss to elephants and help them avoid farming-related debts from financial institutions that otherwise amplify their frustration. The main challenge that might arise with improving government response to HEC is the possibility that it could encourage cropland expansion which may further affect elephants’ habitats. Therefore, the implementation of such strategy needs to go hand in hand with proper land use planning to protect and restore elephants migration corridors and dispersal areas. The importance of this approach cannot be overstated as cropland expansion and conversion of natural habitats are the main drivers for increasing HEC; therefore, land use planning at a landscape scale to secure migration corridors between protected areas is critical. Land use/land cover analysis confirmed that cropland has expanded during the past three decades, bringing humans and elephants into closer contacts. This expansion is agreeably the cause of increasing HEC, and therefore controlling it is critical in improving coexistence between people and elephants. While it may not be practical to build a fence along corridors from Mikumi and Nyerere national parks to Wami-Mbiki WMA due to their great lengths, considerations to reallocate crop fields from within the corridor areas to other areas with comparable agricultural potential should be made to reduce negative interactions.

Similarly, conservation authorities should revisit their benefit sharing plans and allocate more tangible benefits to the communities. The benefits should take into consideration direct costs incurred from HEC and focus on offsetting them. Although constructing classrooms and providing study desks count towards investment in education and long-term development, the immediate needs and challenges associated with HEC should also be met and addressed. As rightfully asked by a woman in Kanyenja “*What is the use of building schools if our kids cannot attend due to fear of elephants…we do not see that as a benefit!”* Elderly people are particularly affected since they have no one to help them, and they do not perceive building classrooms as of any help for their current challenges in dealing with HEC. Furthermore, upgrading part of Selous Game Reserve to Nyerere National Park should be used as the opportunity to bring more benefits to the local communities in the form of payments directly provided as social fund.

Moreover, the national park status presents an opportunity for the local communities to diversify their livelihood activities. Through its Community Conservation Services (CCS) Outreach Program, TANAPA could leverage on expanded tourism opportunity and create conducive environment for camping sites, hospitality industry and provision of related goods and services. Having alternative income-generating activities will reduce over-dependence on subsistence farming and supplement the income, helping to reduce local food insecurity. In Karatu and Mto wa Mbu in northern Tanzania, for example, tourism has, to some extent, helped local communities offset the costs of living with elephants. One opportunity that exists to realize livelihood strategies diversification is a project under implementation by TANAPA and the Ministry of Natural Resources and Tourism, known as Resilient Natural Resource for Tourism and Growth (REGROW), a World Bank funded project aimed at transforming tourism sector in the southern tourism circuit (49,(50)). Specifically, REGROW targets to improve accessibility of the protected areas in southern Tanzania through air or surface transport, improve economic opportunities and diversify tourism products to benefit the local communities(51). Successful implementation of this project will not only contribute to improving livelihoods of the local communities, but also provide enough resources to improve the capacity of the wildlife authorities to respond to HEC.

## Acknowledgements

We acknowledge financial support received from the Center for Landscape Dynamics, Africana Research Center and the Department of Geography at Pennsylvania State University, as well as from the Hamer Foundation. We also thank TAWIRI, TAWA, and the Commission for Science and Technology (COSTECH) for research permits. Members of Tanzania Elephant Foundation (TEF) assisting in data collection: Lameck Mkuburo, Mathayo Ramadhan, Violeth Sanga and Alpha Mfilinge.

## Author Contributions

Conceptualization: Grace S. Malley

Formal Analysis: Grace S. Malley

Funding Acquisition: Grace S. Malley

Methodology: Grace S. Malley; L.J. Gorenflo.

Supervision: L.J. Gorenflo

Writing original draft: Grace S. Malley

Writing-review and editing: Grace S. Malley, L.J. Gorenflo.

## Supporting Information

**S1 table:** Total population and sample sizes from each of the selected village in the study area.

**S2 file**: IRB Clearence

**S3 table:** The number of community members who participated focus group discussion in each village and district.

**S4 table:** Summary of household survey for the study area and for each district.

**S5 file:** Household questionnaire

**S6 file:** Discussion guide

**S7:** Research permit

